# Anticodon sequence determines the impact of mistranslating tRNA^Ala^ variants

**DOI:** 10.1101/2022.11.23.517754

**Authors:** Ecaterina Cozma, Megha Rao, Madison Dusick, Julie Genereaux, Ricard A. Rodriguez-Mias, Judit Villén, Christopher J. Brandl, Matthew D. Berg

## Abstract

Transfer RNAs (tRNAs) maintain translation fidelity through accurate charging by their cognate aminoacyl-tRNA synthetase and codon:anticodon base pairing with the mRNA at the ribosome. Mistranslation occurs when an amino acid not specified by the genetic message is incorporated into proteins and has applications in biotechnology, therapeutics and is relevant to disease. Since the alanyl-tRNA synthetase uniquely recognizes a G3:U70 base pair in tRNA^Ala^ and the anticodon plays no role in charging, tRNA^Ala^ variants with anticodon mutations have the potential to mis-incorporate alanine. Here, we characterize the impact of the 60 non-alanine tRNA^Ala^ anticodon variants on the growth of *Saccharomyces cerevisiae*. Overall, 36 tRNA^Ala^ anticodon variants decreased growth in single-or multi-copy. Mass spectrometry analysis of the cellular proteome revealed that 52 of 57 anticodon variants, not decoding alanine or stop codons, induced mistranslation when on single-copy plasmids. Variants with G/C rich anticodons resulted in larger growth deficits than A/U rich variants. In most instances, synonymous anticodon variants impact growth differently, with anticodons containing U at base 34 being the least impactful. For anticodons generating the same amino acid substitution, reduced growth generally correlated with the abundance of detected mistranslation events. Differences in decoding specificity, even between synonymous anticodons, resulted in each tRNA^Ala^ variant mistranslating unique sets of peptides and proteins. We suggest that these differences in decoding specificity are also important in determining the impact of tRNA^Ala^ anticodon variants.

## Introduction

Transfer RNAs (tRNAs) are adaptor molecules that decode the genetic message contained within mRNA into protein sequence during translation. Translation fidelity and proteome accuracy is maintained by tRNAs at two steps. First, tRNAs are recognized by their aminoacyl-tRNA synthetase (aaRS) and charged with the cognate amino acid. Second, accurate base pairing between the tRNA anticodon and mRNA codon during decoding at the ribosome ensures the correct amino acid is incorporated into the growing polypeptide chain [1–3]. Errors at either step lead to mistranslation, the incorporation of an amino acid not specified by the genetic message. Mis-incorporation events occur naturally in cells with estimated frequencies of 10^−5^ to 10^−3^ depending on the codon [4–7]. In addition, mistranslation increases in response to environmental conditions [8–10] and in the presence of tRNA variants [11–14].

Aminoacyl-tRNA synthetases recognize their cognate tRNAs through individual bases or structural motifs within the tRNA called identity elements [15–17]. For most tRNAs, the anticodon is the main identity element, providing a direct link between the genetic code and amino acid being incorporated. This is not the case for tRNA^Ala^ and tRNA^Ser^, which are recognized through identity elements outside of the anticodon [18–21]. Specifically, the alanyl-tRNA synthetase (AlaRS) recognizes a conserved G3:U70 base pair in the tRNA^Ala^ acceptor stem (Figure 1A) [18,19]. Seryl-tRNA synthetase recognizes a uniquely long variable arm 3’ of the anticodon stem in tRNA^Ser^ [20,21]. Identity elements for leucylation of tRNA^Leu^ is species specific, but is also largely independent of the anticodon [22–24]. Consequently, with one or two mutations to the anticodon tRNA^Ala^, tRNA^Ser^ and tRNA^Leu^ are theoretically capable of mis-incorporating alanine, serine or leucine, respectively, at non-cognate codons [13,14,25,26].

**Figure 1.**
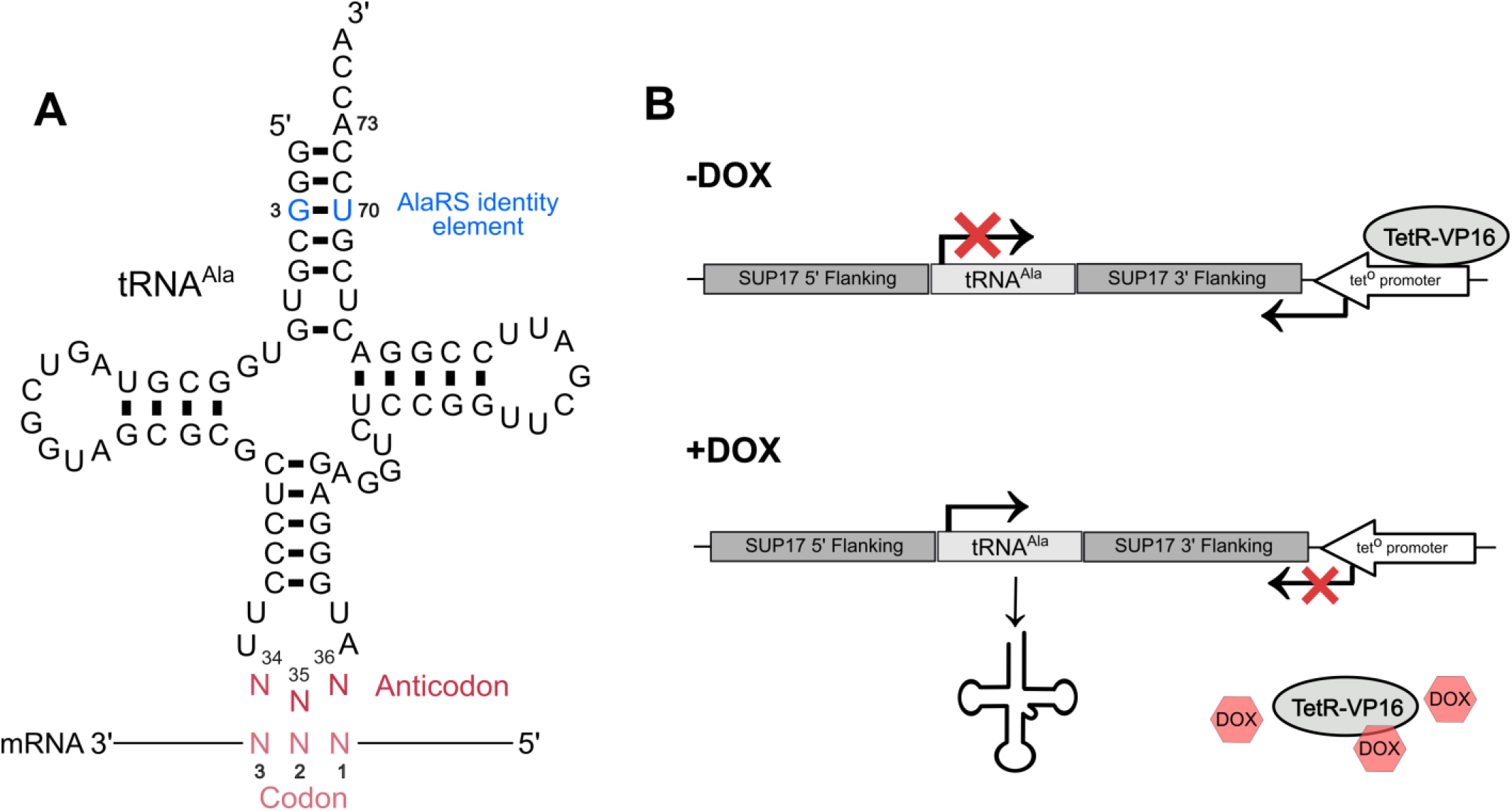
Schematic of tRNA^Ala^ containing a degenerate anticodon and the tetracycline inducible tRNA expression system. **(A)** The structure of tRNA^Ala^ with a degenerate anticodon. Bases colored in blue represent the G3:U70 base pair required for recognition and charging by AlaRS. The anticodon is shown in red. **(B)** tRNA^Ala^ in the tetracycline inducible system used to regulate tRNA^Ala^ anticodon variant expression flanked by up- and downstream *SUP17* sequence. In the absence of doxycycline, the tet^O^ promoter is bound by the TetR-VP16 transcriptional activator which represses tRNA expression by driving RNA polymerase II expression across the tRNA gene. In the presence of doxycycline, TetR-VP16 binds doxycycline and dissociates from the promoter allowing the tRNA to be transcribed by RNA polymerase III.

When introduced into cells, tRNA variants capable of mistranslation give rise to a heterogeneous mixture of molecules arising from the same genetic template but differing at a subset of amino acid positions. Carl Woese described these as statistical proteins which can provide cells with a wider range of functions than would be achievable from a homogeneous proteome [27]. As further indicated by Woese, statistical proteins form the basis of a powerful strategy for searching protein phase space to find novel protein activities [28].

Mistranslating tRNA variants have implications for health and disease. Santos *et al*. demonstrated that mistranslating tRNAs induce cell transformation and result in faster growing tumours in mice [12]. Flies and nematodes expressing mistranslating tRNAs show morphological defects [29,30]. Mistranslating flies also have locomotive defects similar to those seen for flies expressing alleles associated with neurodegeneration [29,31–33]. Consistent with a possible role in disease, tRNA variants with the potential to mistranslate are found in human populations [34,35]. Lant *et al*. demonstrated that in cell models of neurodegeneration a naturally occurring mistranslating tRNA variant slowed protein aggregate formation and inhibited the clearance of protein aggregates [36]. Further to their impact on health and disease, mistranslating tRNAs have potential in gene therapy as first demonstrated for β-thalasemia by Temple *et al*. [37]. Improvements in methods to introduce genes and RNA into cells have resulted in a renewed exploration of suppressor tRNA therapies being developed to treat diseases arising from premature termination codons, such as Duchenne muscular dystrophy and cystic fibrosis [38,39]. Lastly, mistranslating tRNA variants provide novel tools for determining the effects of thousands of missense mutations simultaneously [40], enabling the functional interpretation of missense mutations at a genome-wide level for personalized medicine applications.

The diverse applications and disease implications of mistranslating tRNAs led us to engineer all possible anticodon variants of *Saccharomyces cerevisiae* tRNA^Ala^ and measure their impact on growth and ability to induce mistranslation. We hypothesized that the non-alanine anticodon variants would impact cells differently, as a result of making different amino acid substitutions. Indeed, differences in growth inhibition and mistranslation were observed for both non-synonymous and synonymous anticodons when found in the context of tRNA^Ala^. Impact was generally greatest for G/C rich anticodons and less for anticodons with U at base 34. The abundance of mistranslated peptides caused by the tRNA^Ala^ variant, the specific proteins and positions mistranslated and the type of amino acid substitution contributed to the extent cell growth was affected by each variant.

## Materials and Methods

### Yeast strains and growth

Wild type haploid yeast strains are derivatives of BY4742 (***MAT****α his3Δ1 leu2Δ0 lys2Δ0 ura3Δ0*) [41]. The haploid strain CY8652 (***MAT****α his3Δ1 leu2Δ0 lys2Δ0 ura3Δ0 tTA*-URA3)* containing the tet “off” activator (tTA*) marked with *URA3* was derived from R1158 [42] after crossing with BY4741 and sporulation.

Yeast strains were grown at 30°C in yeast peptone medium or in synthetic medium supplemented with nitrogenous bases and amino acids containing 2% glucose. Transformations were performed using the lithium acetate method as previously described [43].

### Growth assays

*Inducible tRNA*^*Ala*^ *anticodon variants*. Yeast strain CY8652 constitutively expressing the TetR-VP16 protein and containing a *LEU2* plasmid expressing either a control tRNA^Ala^ _GGC(Ala)_ decoding alanine codons or the anticodon variants were grown to saturation in medium lacking uracil and leucine without doxycycline. Strains were diluted to OD_600_ of 0.1 in the same medium containing 0, 0.01 or

1.0 µg/mL of doxycycline and grown for 24 hours at 30°C with agitation. OD_600_ was measured every 15 minutes for 24 hours in a BioTek Epoch 2 microplate spectrophotometer. Doubling time was calculated using the R package “growthcurver” [44] and was normalized to the wild-type strain grown at the respective doxycycline concentration. Each strain was assayed in triplicate.

*2*μ *tRNA*^*Ala*^ *anticodon variant derivatives*. Yeast strain BY4742 was transformed with 1.0 μg of plasmid expressing the control tRNA^Ala^ _CGC(Ala)_ variant or anticodon variant tRNAs and plated onto minimal medium lacking uracil. Plates were imaged after 2 days of growth and quantified using the ImageJ [45] “Watershed” package.

### DNA constructs

The construct encoding the tRNA^Ala^ variants was synthesized as a GeneString fragment (ThermoFisher) containing a random mix of each nucleotide at anticodon positions 34, 35 and 36 and the tRNA^Ala^ gene was flanked by approximately 300 base pairs of up- and downstream sequence from *SUP17*. The synthetic gene fragment was cloned as a *Hin*dIII/*Not*I fragment into pCB4699, a *LEU2* centromeric plasmid (YCplac111) containing a tetracycline regulated promoter downstream of the tRNA gene as previously described [46]. For variants that were not selected out of the randomized anticodon pool, constructs expressing these tRNA^Ala^ variants were engineered via two-step PCR reactions. To reduce the number of primers needed, mismatched primer sets were used to generate two variants from a single PCR reaction. PCR fragments were subcloned into pGEM-Teasy (Promega) and then moved as *Hin*dIII/*Not*I fragments into pCB4699. The sequences of the 64 tRNA^Ala^ anticodon variants generated and plasmid numbers are listed in Table S1. The cloning approach and primers used to generate each variant (if applicable) are listed in Table S2.

Multicopy plasmids expressing tRNA^Ala^ variants were made by cloning *Hin*dIII/*Eco*RI fragments containing each tRNA variant into YEplac195. Plasmid numbers are listed in Table S3.

### Mass spectrometry

Strains were grown overnight, in quadruplicate, in medium lacking leucine and uracil. Cultures were diluted to OD_600_ of 0.01 in the same medium containing 1.0 µg/mL doxycycline except for the strain expressing tRNA^Ala^ _CGG(Pro)_ which was diluted to OD_600_ of 0.1. Cultures were harvested by washing once in 1x yeast nitrogen base and frozen in liquid nitrogen when they reached an OD_600_ between 0.8–1.0, except for tRNA^Ala^ _GCC(Gly)_ which due to its slow growth was harvested 36 hours after dilution at OD_600_ ~ 0.4.

Cells were lysed by bead-beating with 0.5 mm glass beads at 4°C in urea lysis buffer (8 M urea, 50 mM Tris pH 8.2, 75 mM NaCl). Lysates were cleared by centrifugation at 21,000 x g for 10 minutes at 4°C and protein concentration was determined by Bicinchoninic acid assay (Pierce, ThermoFisher Scientific). Proteins were reduced with 5 mM dithiothreitol for 30 minutes at 55°C, alkylated with 15 mM iodoacetamide for 30 minutes at room temperature in the dark and the alkylation was quenched with an additional 5 mM dithiothreitol for 30 minutes at room temperature. For each sample, 50 µg of protein was diluted 4-fold with 50 mM Tris pH 8.9 and digested for 4 hours at 37°C with 1.0 µg LysC (Wako Chemicals). Digestions were acidified to pH 2 with trifluoroacetic acid and desalted over Empore C18 stage tips [47].

For strains containing the variants with UUU(Lys) and CUU(Lys) anticodons and the corresponding control strain containing tRNA^Ala^ _GCC(Ala)_ samples were lysed as above in 4 M guanidine hydrochloride, 100 mM triethylammonium bicarbonate pH 8.0. Samples were prepared as above, except after quenching the alkylation reaction, 100 µg of protein was acetylated with 200 µg of Sulfo-NHS-acetate (Pierce, ThermoFisher Scientific) for 1 hour at room temperature. Robotic purification and digestion of proteins into peptides was performed on the Kingfisher Flex (ThermoFisher Scientific) using trypsin and the R2-P1 method [48]. Digestions were acidified and desalted as above. Peptides were resuspended in 200 mM EPPS pH 9.5 with 5% hydroxylamine and incubated for 5 hours at room temperature to reduce acetylation on serine, threonine and tyrosine residues before peptides were desalted again.

Peptide samples were resuspended in 4% acetonitrile, 3% formic acid and subjected to liquid chromatography coupled to tandem mass spectrometry on a tribrid quadrupole orbitrap mass spectrometer (Orbitrap Eclipse; ThermoFisher Scientific). Samples were loaded onto a 100 µm ID x 3 cm precolumn packed with Reprosil C18 3 µm beads (Dr. Maisch GmbH) and separated by reverse-phase chromatography on a 100 µm ID x 30 cm analytical column packed with Reprosil C18 1.9 µm beads (Dr. Maisch GmbH) housed in a column heater set at 50°C. Peptides were separated using a gradient of 7-50% acetonitrile in 0.125% formic acid delivered at 400 nL/min over 95 minutes, with a total 120 minute acquisition time. The mass spectrometer was operated in data-dependent acquisition mode with a defined cycle time of 3 seconds. For each cycle, one full mass spectrometry scan was acquired from 350 to 1200 m/z at 120,000 resolution with a fill target of 3E6 ions and automated calculation of injection time. The most abundant ions from the full MS scan were selected for fragmentation using 1.6 m/z precursor isolation window and beam-type collisional-activation dissociation (HCD) with 30% normalized collision energy. MS/MS spectra were acquired at 30,000 resolution by setting the AGC target to standard and injection time to automated mode. Fragmented precursors were dynamically excluded from selection for 60 seconds.

MS/MS spectra were searched against the *S. cerevisiae* protein sequence database (downloaded from the Saccharomyces Genome Database resource in 2014) using Comet (release 2015.01) [49]. The precursor mass tolerance was set to 20 ppm. Constant modification of cysteine carbamidomethylation (57.0125 Da) and variable modification of methionine oxidation (15.9949 Da) was used for all searches. For samples containing the UUU(Lys) and CUU(Lys) anticodon variants and the corresponding control samples, variable modification of lysine acetylation (42.0105 Da) was included. A variable modification of each amino acid to alanine was used for the respective searches. A maximum of 3 of each variable modification was allowed per peptide. Search results were filtered to a 0.05% false discovery rate at the peptide spectrum match level using Percolator [50]. Peptide intensity was determined using in-house quantification software to extract MS1 intensity. To determine if variants were inducing mistranslation, the proportion of identified mistranslated peptides was calculated from the number of unique mistranslated peptides for which the non-mistranslated sibling peptide was also observed. The mistranslated proportion is defined as the number of unique mistranslated peptides, where alanine had been mis-incorporated, divided by the number of total peptides containing the wild-type amino acid and expressed as a percentage. At the codon level, the decoding specificity of each tRNA^Ala^ variant was determined using a custom Perl script (Supplemental File 1) to map codons back to mistranslated residues. Only peptides with one possible substitution event were considered to allow for accurate localization of the mistranslated residue.

The raw mass spectrometry data have been deposited to the ProteomeXchange Consortium via the PRIDE [51] partner repository with the dataset identifier PXD038242 and an annotated list of the file names can be found in Table S4.

### Factors used to correlate with growth impact of each variant

The following factors were correlated using multiple linear regression with the relative growth impact of each variant as the dependent variable: the mistranslation abundance calculated from the mass spectrometry data and the difference of the replaced amino acid relative to alanine for the following five properties: volume [52], electron-ion interaction potential [53], hydrophobicity [54], α-helix propensity [54] and π-helix propensity [55].

### Statistical analyses

Statistical analyses were performed using R studio 1.4.1717. Example code can be found in Supplemental File 2. Welch’s *t*-test was used to compare doubling times of strains containing tRNA^Ala^ anticodon variants to the control tRNA^Ala^ _GGC(Ala)_ strain and corrected for multiple tests using the Bonferroni method. Wilcoxon rank sum test was used to compare median colony size of strains transformed with tRNA^Ala^ anticodon variants on multicopy plasmids relative to the control tRNA^Ala^ _CGC(Ala)_ and corrected for multiple tests using the Bonferroni method. Welch’s *t*-test was used to compare the proportion of mistranslated peptides detected measured for strains containing tRNA^Ala^ anticodon variants to the control tRNA^Ala^ _GGC(Ala)_ strain and corrected for multiple tests using the Benjamini-Hochberg method. For the multiple linear regression, independent variables with *p-* values less than 0.05 after Bonferroni multiple test correction were considered to contribute to the impact each tRNA^Ala^ anticodon variant had on growth. Association between growth impact and estimated mistranslation frequency was determined using Spearman’s correlation.

## Results

### Engineering and characterizing the growth impact of tRNA^Ala^ anticodon variants

To engineer tRNA^Ala^ anticodon variants with the potential to mis-incorporate alanine at non-alanine codons, we cloned tRNA^Ala^ with each of the 64 possible anticodon sequences into a tetracycline inducible system [46] to regulate tRNA expression and control mistranslation-associated toxicity (Figure 1, Table S1). In the absence of the tetracycline analog doxycycline, tRNA expression is repressed. When doxycycline is added to the growth medium, the tRNA variant is expressed in a titratable manner. Since we previously found that regulation of this system was improved with tRNA^Ser^ *SUP17* flanking sequence [46], the tRNA^Ala^ encoding gene was positioned precisely into the *SUP17* locus (Figure S1). As such, all anticodons were examined in the same context and of note, the anticodon should not play a role in the stability or expression of a tRNA [13].

To assess the impact of the tRNA^Ala^ variants on yeast growth, plasmids containing one of 64 tRNA^Ala^ anticodon variants were transformed into a yeast strain constitutively expressing TetR-VP16 (CY8652) [46]. Transformants were isolated and their growth characterized in liquid minimal media containing 0, 0.01 or 1.0 µg/mL doxycycline. Growth curves were performed in biological triplicate for each strain at 30°C over 24 hours. Figure 2A shows representative growth curves for strains containing plasmids with the control tRNA^Ala^ _GGC(Ala)_ or tRNA^Ala^ variants with CAU(Met), GAA(Phe) and AGG(Pro) anticodons as examples of those having minimal, intermediate and high impact on growth, respectively. As expected, impact on growth was less severe at lower doxycycline concentrations for tRNA^Ala^ variants with GAA(Phe) and AGG(Pro) anticodons. Growth of strains with the control tRNA^Ala^ _GGC(Ala)_ or the variant with a CAU(Met) anticodon was unchanged at all doxycycline concentrations.

**Figure 2.**
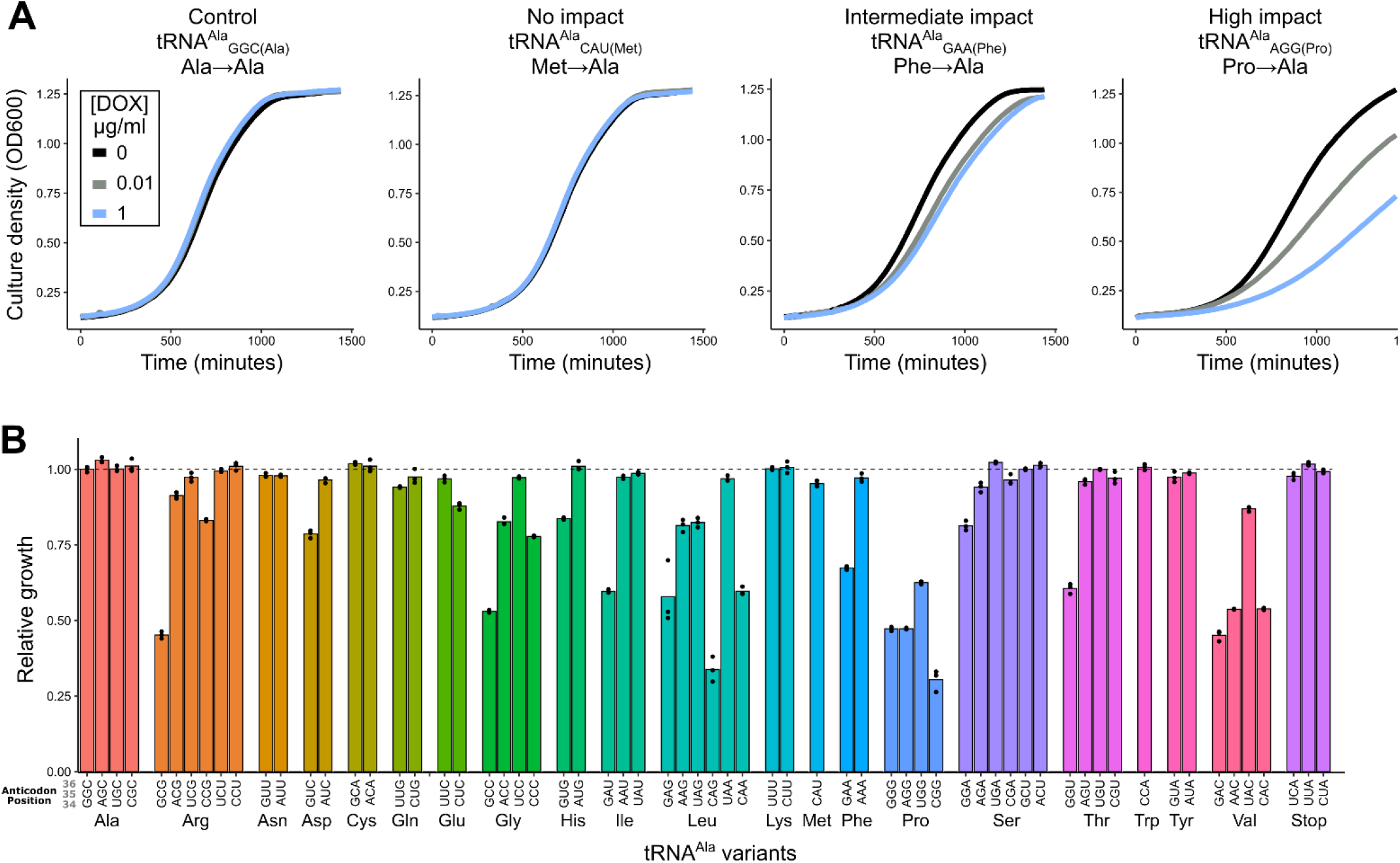
Impact of tRNA^Ala^ anticodon variants on yeast growth. **(A)** Representative growth curves of yeast strain CY8652 containing a plasmid with the control tRNA^Ala^ _GGC(Ala)_, a tRNA^Ala^ _CAU(Met)_ variant with no growth impact, a tRNA^Ala^ _GAA(Phe)_ variant with intermediate growth impact or a tRNA^Ala^ _AGG(Pro)_ variant with a severe growth impact. Black, grey and blue lines represent strains grown in media containing 0, 0.01 and 1.0 μg/mL doxycycline, respectively. **(B)** Yeast strain CY8652 containing plasmids with the control tRNA^Ala^ _GGC(Ala)_ or one of the tRNA^Ala^ anticodon variants were grown for 48 hours at 30°C in medium lacking uracil and leucine, diluted to an OD_600_ of 0.1 in the same medium with 1.0 μg/mL doxycycline and grown for 24 hours at 30°C with agitation. OD_600_ was measured at 15-minute intervals and doubling time was quantified. Relative growth of tRNA^Ala^ anticodon variants was calculated by multiplying the inverse of each variant doubling time by the average doubling time of the tRNA^Ala^ _GGC(Ala)_ control (for raw data and statistical comparisons see Table S5 and S6). Each point represents one biological replicate (n = 3).

The relative growth of each strain containing one of 64 possible tRNA^Ala^ variants in medium with 1.0 µg/mL doxycycline is shown in Figure 2B and relative growth at all three doxycycline concentrations is shown in Figure S2 (raw data can be found in Table S5). Of the 60 variants with non-alanine anticodons, 25 significantly decreased growth compared to the control tRNA^Ala^ _GGC(Ala)_ (Table S6). We observed a range of relative growth rates indicating that tRNA^Ala^ variants with anticodons decoding the 19 non-alanine amino acids have diverse impacts. For example, G/C rich variants with CGG(Pro), GAC(Val), GCC(Gly) and CAG(Leu) anticodons resulted in the greatest growth reduction. A moderate growth reduction was caused by variants with GAA(Phe), GUG(His) and GGA(Ser) anticodons. Anticodon variants that caused no reduction in growth included the A/U rich variants with AAA(Phe), AAU(Ile), UAU(Ile), UUU(Lys), and AUU(Asn) anticodons as well as variants with anticodons decoding the three stop codons. The minimal impact of anticodon variants decoding stop codons likely reflects strong competition from release factors. Interestingly, different growth impacts were observed for variants decoding synonymous codons. For example, the variant with a GGU(Thr) anticodon reduced growth to 61% of the control tRNA^Ala^, whereas the other three threonine decoding variants had a minimal effect.

To analyze the importance of anticodon G/C content on the impact of tRNA^Ala^ variants, we plotted the relative growth of strains containing each variant on Grosjean and Westhof’s alternative representation of the genetic code [56], which places ‘strong’ G/C rich anticodons at the top of the circle plot and ‘weak’ A/U rich anticodons at the bottom as determined by the Turner binding energy of the base pairing between the first and second positions of the codon with bases 35 and 36 of the anticodon. As shown in Figure 3A, ‘strong’ anticodons generally resulted in a greater reduction in growth compared to ‘weak’ anticodons.

**Figure 3.**
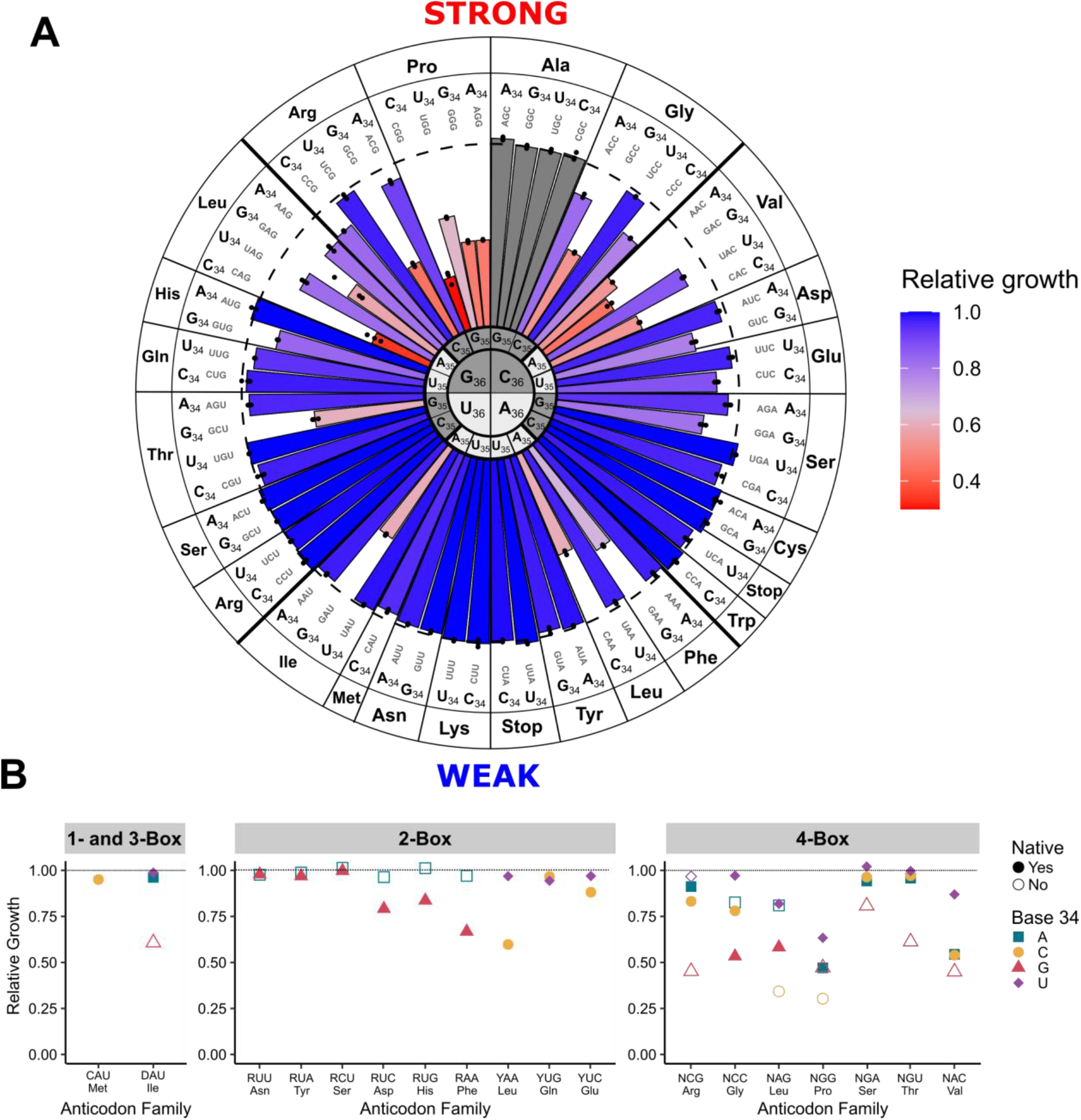
Anticodon sequence affects growth impact. **(A)** Relative growth for each tRNA^Ala^ variant as in Figure 2B plotted for each anticodon on Grosjean and Westhof’s alternative representation of the genetic code [56] where ‘strong’ codon:anticodon pairs are at the top of the circle plot and ‘weak’ pairs are at the bottom. The length of each bar is proportional to the relative growth of each variant. **(B)** Relative growth for mistranslating tRNA^Ala^ anticodon variants decoding synonymous anticodons differing at base 34. Variants are grouped by number of different possible anticodons decoding a specific amino acid (1-, 2-, 3-or 4-box) with the same base at positions 35 and 36. Amino acid identity is listed below the anticodon. Native and non-native anticodon sequences in yeast are depicted with filled and empty shapes, respectively. D: adenine, guanine or uracil; R: purine; Y: pyrimidine; N: any base.

Next, we evaluated how the identity of tRNA base 34, the first position of the anticodon, impacts growth. Figure 3B shows the relative growth of strains containing synonymous anticodon variants decoding the same amino acid and differing only at base 34. In four-box sets, variants with U at position 34 had less impact on growth as compared to their A, G or C counterparts. Similarly, two-box variants with A at position 34 had less impact on growth than their G-containing counterparts. Consistently, G or C at position 34 had the greatest impact on growth in each isodecoder set.

Of the 57 possible anticodons not decoding alanine or stop codons, 18 are not natively found in *S. cerevisiae*. In Figure 3B, non-native and native anticodons are represented by open and filled points, respectively. For the three- and four-box tRNA variants including those decoding Ser, Arg and Leu codons, the most impactful anticodon is non-native in seven of eight cases (the exception being the variant with GCC(Gly) anticodon). This pattern does not hold in any of the three cases for the two-box tRNAs where it can be evaluated.

### Increasing tRNA^Ala^ anticodon variant copy number reveals growth impact

In single copy, 35 non-alanine variants did not statistically impact growth when compared to the control strain expressing tRNA^Ala^ _GGC(Ala)_. Since each tRNA^Ala^ anticodon variant competes with the native tRNA species for decoding, we cloned a subset of the variants into a 2*µ* multicopy plasmid to determine if increasing copy number would produce a growth phenotype. The wild-type BY4742 strain was transformed with the 2*µ* plasmids and transformant colony size quantified relative to a control transformation of tRNA^Ala^ _CGC(Ala)_ on a multicopy plasmid (Figure 4, Figure S3, Table S7). selection pressure of deleterious plasmids towards lower copy. As a control to demonstrate that overexpressing a mistranslating variant exacerbates growth impact, we also analyzed the GAA(Phe) anticodon variant that resulted in slow growth on a single-copy plasmid. When this variant was expressed on a 2*µ* plasmid, no transformants were obtained. Of the 17 low impact variants tested, 12 significantly decreased median colony size compared to the control when introduced on a multicopy plasmid. Four of these led to mean colony size 50% less than the control (variants with UCC(Gly), CAU(Met), AGU(Thr), AAA(Phe) anticodons) and one variant produced transformants that were too small to quantify (variant with CUG(Gln) anticodon). These variants likely mistranslate in single copy, but at a low frequency that does not lead to a growth phenotype. In contrast, five tRNA^Ala^ variants showed no statistical impact on colony size, even in high copy number (variants with GCA(Cys), UUU(Lys), CCA(Trp), AUA(Tyr), and CUA(Stop) anticodons).

**Figure 4.**
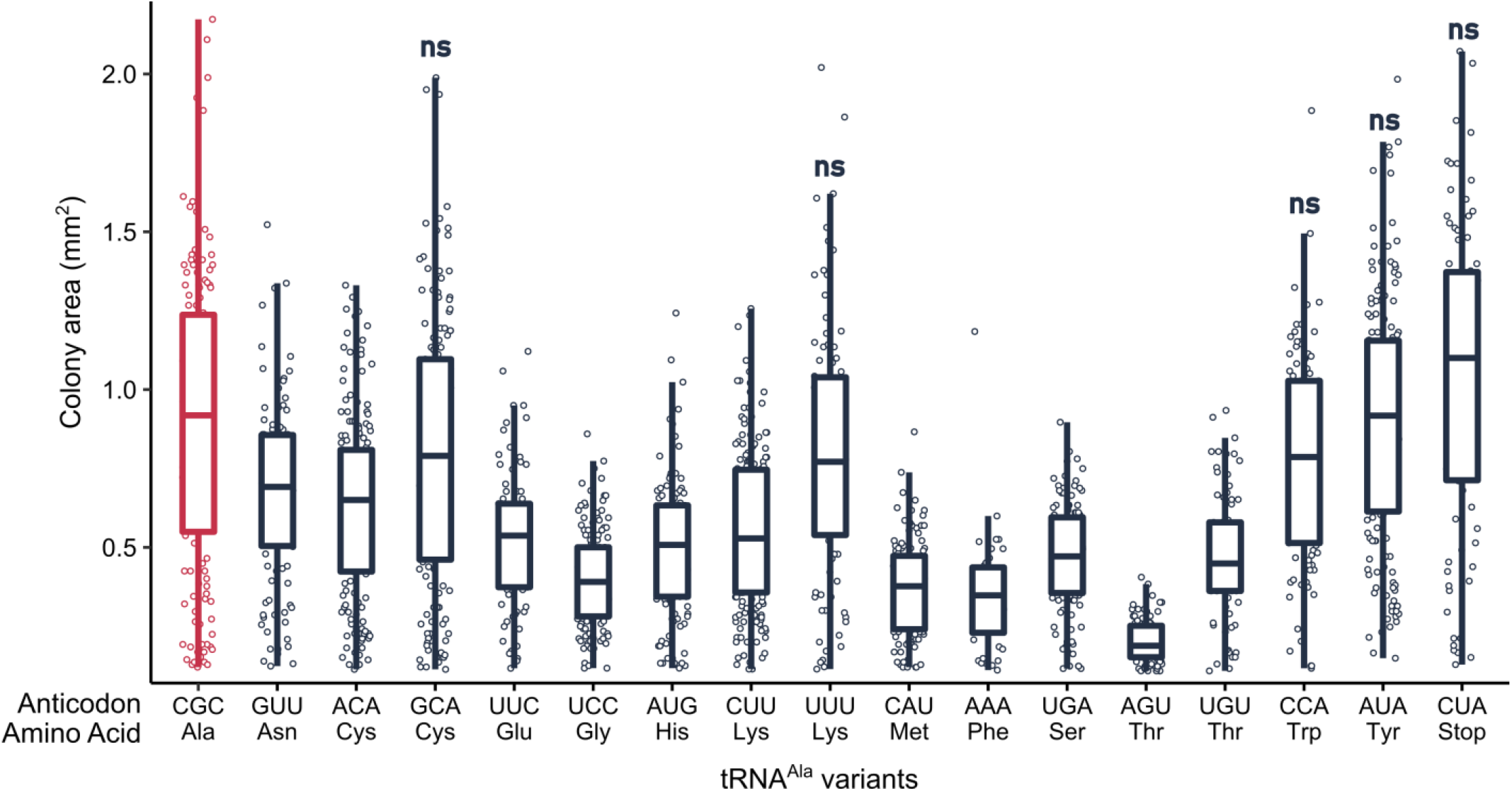
Overexpressing some but not all tRNA^Ala^ variants reduce growth. Strain BY4742 was transformed with 1.0 µg of a *URA3* multicopy plasmid containing each tRNA^Ala^ anticodon variant, plated on media lacking uracil and grown at 30°C. Transformants were imaged after 48 hours (raw images are in Figure S3) and colony area was quantified using the ImageJ ‘Watershed’ package. Each point represents one colony and horizontal bars at the center of each boxplot represent median colony size. Median colony size of each variant was compared to the control tRNA^Ala^ _CGC(Ala)_ (shown in red) using a Wilcoxon rank sum test with Bonferroni correction to determine significance (see Table S7 for *p*-values). Variants lead to a statistically significant decrease in growth (*p* < 0.01) unless denoted by ‘ns’.

### Estimating mistranslation frequency of each tRNA^Ala^ anticodon variants

To evaluate the relationship between reduced growth caused by the tRNA^Ala^ variants and their ability to induce mistranslation, we analyzed the cellular proteome by mass spectrometry after growth of each strain in medium containing 1.0 μg/mL doxycycline and isolation of cellular protein in denaturing conditions. To determine if mistranslation was occurring, we calculated the fraction of unique mistranslated peptides relative to the total number of peptides containing the wild-type amino acid (Figure 5A, Table S8). Of the 57 non-alanine anticodon variants measured, 52 resulted in statistically elevated levels of mistranslated peptides compared to the control strain (variants decoding stop codons were not analyzed). Similar to the effect of these variants on growth, the proportion of mistranslated peptides detected spanned a wide range. For example, we detected 12.8%, 11.9% and 11.6% of peptides mistranslated for variants UGG(Pro), GAC(Val) and GUG(His) anticodons, respectively, whereas we detected 0.12% of peptides mistranslated for the variant with an AAA(Phe) anticodon. It is important to recognize that these values are estimates, as the measured proportion of peptides mistranslated reflects specific properties of the mistranslated versus wild-type peptides, such as differences in protein turnover and the ability to detect individual peptides.

**Figure 5.**
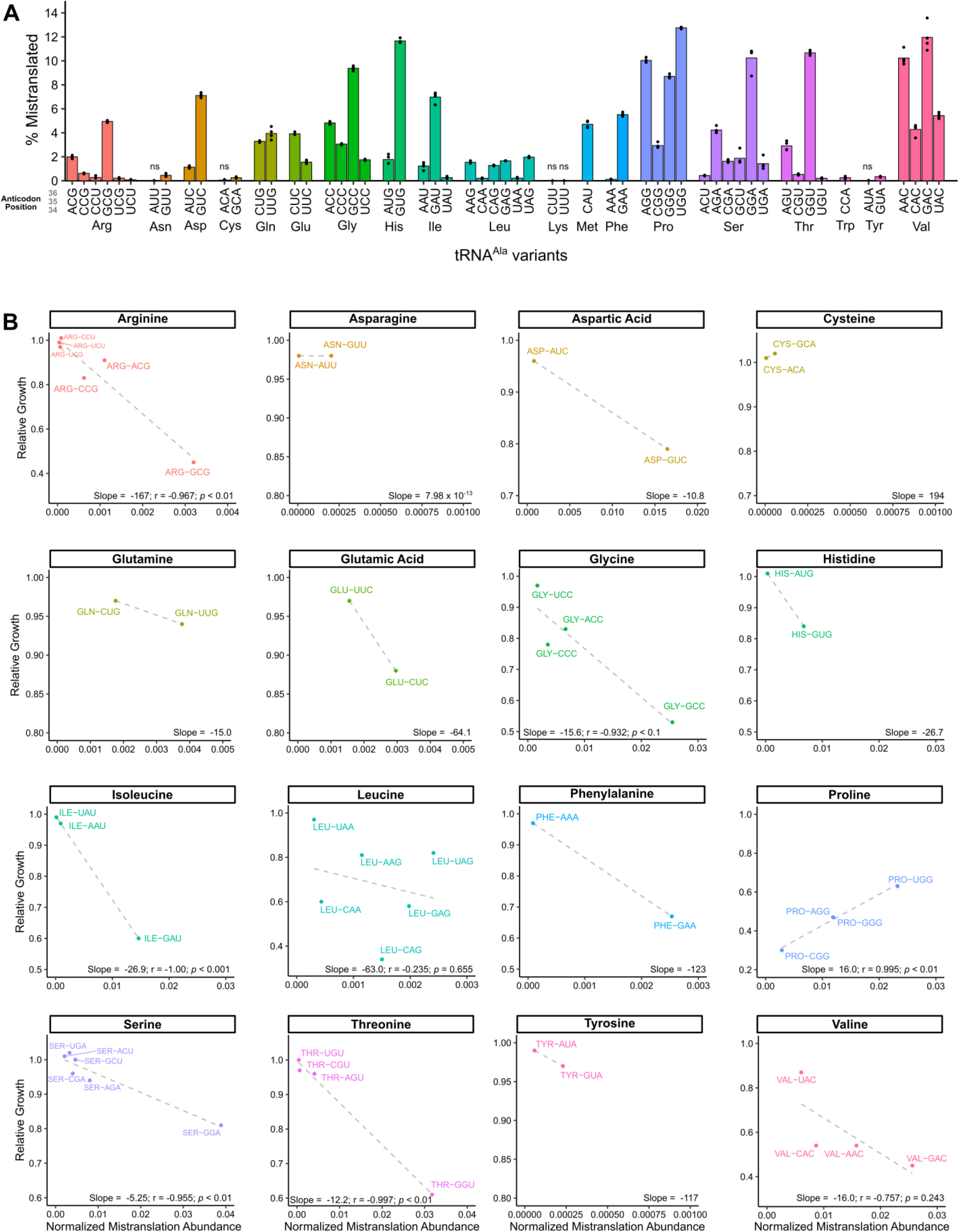
Mistranslation detected by mass spectrometry-based analysis of the cellular proteome for strains containing tRNA^Ala^ anticodon variants. **(A)** Yeast strain CY8652 containing a tRNA^Ala^ anticodon variant were diluted into medium containing 1.0 μg/mL doxycycline to induce tRNA expression and harvested at an OD_600_ of ~ 1.0. Cells were lysed in a denaturing buffer and mass spectrometry analysis of the cellular proteome was performed. The proportion of peptides detected as mistranslated, expressed as a percentage, was calculated for each variant from the number of unique peptides where alanine mis-incorporation was observed relative to the number of unique peptides where the wild-type amino acid was observed. Each point represents one biological replicate (n ≥ 3). Variants have statistically elevated levels of mistranslated peptides relative to the control strain containing tRNA^Ala^ _GGC_ (Welch’s *t*-test; Benjamini-Hochberg corrected *p* < 0.01) unless denoted by ‘ns’. **(B)** Summed MS1 intensity of mistranslated peptides normalized to the summed MS1 intensities of all detected peptides (normalized mistranslation abundance) plotted against the relative growth for each tRNA^Ala^ variant for each amino acid family. Amino acids with one anticodon (Trp and Met) and Lys where neither anticodon variant gave a detectable level of mistranslation are not shown.

We hypothesized that the impact each variant has on growth is influenced by the abundance of mistranslated proteins present in the cell under steady-state conditions, which takes into account how frequently mistranslation occurs and the number of sites mistranslated throughout the proteome. Therefore, we calculated the summed MS1 intensity of detected mistranslated peptides normalized to the summed MS1 intensity of all peptides and plotted this against relative growth (Figure S4). There was a weak but significant correlation, where variants that had a greater overall abundance of mistranslated peptides tended to have more impact on growth (linear *r*^*2*^ = 0.240, Spearman’s rank correlation *p* = 1.81 × 10^−9^).

Since the type of amino acid substitution would also be expected to influence the impact on growth, we plotted the estimated abundance of mistranslated peptides versus relative growth separately for the 19 non-alanine isoacceptor sets (Figure 5B). In general, as the estimated abundance of mistranslated peptides increased, the impact on growth was more severe. Negative correlation coefficients greater than 0.9 were seen for 5 of the 8 cases where the number of anticodons allowed correlation coefficients to be calculated. Indicative of some substitutions being better tolerated than others, the slope of the line reflecting the relationship between estimated total mistranslation and growth differed between amino acid families. The arginine to alanine and serine to alanine substitution plots had the greatest and least slopes, respectively. The presence of additional factors impacting growth in an anticodon specific fashion was apparent from the variants with anticodons decoding glycine, leucine and valine codons where reduced growth caused by one or more variants did not correlate well with the summed MS1 intensity of the mistranslated peptides. In contrast to the other variants, the plot for variants mis-incorporating alanine at proline codons showed that NGG(Pro) anticodon variants causing less mistranslation had a greater impact on growth.

The amino acid specific differences in the slopes of the mistranslation abundance versus relative growth plots support that the chemical properties of the substituted amino acid relative to alanine are a factor in determining the impact of the mistranslation event on growth. To determine if we could attribute this relationship to a specific factor, we performed multiple linear regression using the difference between each replaced amino acid and alanine for five factors identified by Then *et al*. that distinguish the proteinogenic amino acids [57]: (1) volume [52]; (2) electron-ion interaction potential [53]; (3) hydrophobicity [54]; (4) α-helix propensity [54] and (5) π-helix propensity [55]. When these factors were included, the correlation with growth improved (linear *r*^*2*^ = 0.470, *p* = 1.06 × 10^−5^) as compared to when we only considered mistranslation abundance. Volume, electron-ion interaction potential and π-helix propensity significantly contributed to the improved correlation.

### Synonymous anticodons target different proteins and positions

For the variants that substitute glycine, valine and leucine with alanine, reduced growth caused by one or more variants did not correlate well with the summed MS1 intensity of the mistranslated peptides. The tRNAAla variants with leucine anticodons resulted in mistranslation amounts that were the least correlated with their effect on growth (Figure 5B). The UAG(Leu) anticodon variant induced the greatest amount of mistranslation within this set and reduced growth to ~80% of the control strain, whereas the CAG(Leu) variant induced an intermediate amount of mistranslation but reduced growth to less than 40%. We hypothesized that the different impact on growth might be explained by codon specific mistranslation by the tRNA^Ala^ variants, resulting in the mistranslation of different positions within different targeted proteins. Figure 6A shows codons that were mistranslated for the four tRNA^Ala^ variants with NAG leucine anticodons. The decoding specificity for variants with UAG(Leu) and CAG(Leu) anticodons was distinct from each other and from the variants with AAG(Leu) and GAG(Leu) anticodons, which were similar. In line with the differences in decoding specificity, there was minimal overlap in the mistranslated peptides and proteins identified in strains containing the variants with UAG(Leu) and CAG(Leu) anticodons (Figure 6B). In contrast, variants with AAG(Leu) and GAG(Leu) anticodons mistranslated relatively similar sets of peptides and proteins and their growth impacts were proportional to their estimated mistranslation abundance.

**Figure 6.**
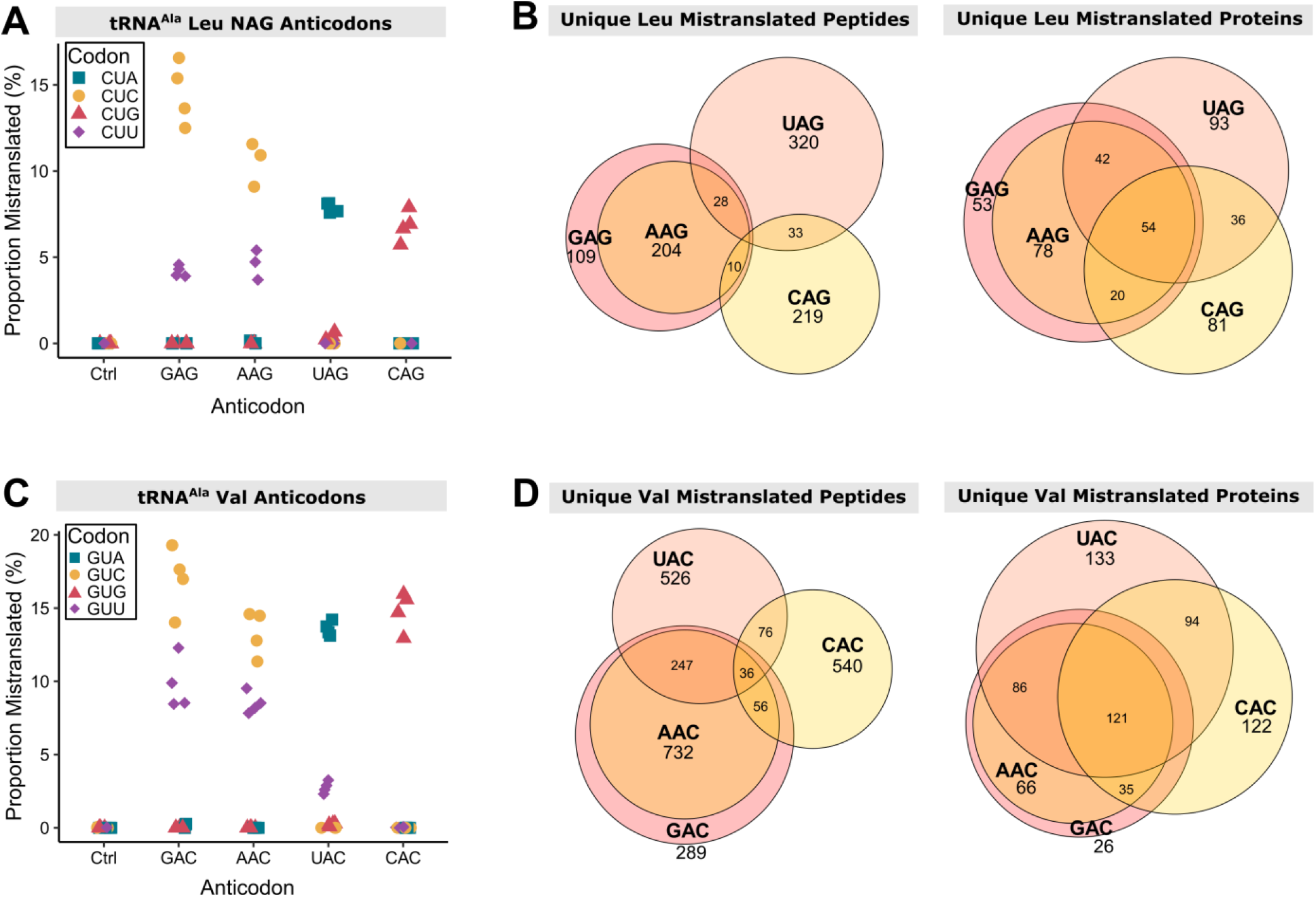
tRNA^Ala^ variants with leucine NAG anticodons or valine NAC anticodons mistranslate at unique codons leading to different subsets of the proteome experiencing mistranslation. **(A)** Codon specific proportion of unique mistranslated peptides for the control strain containing tRNA^Ala^ _GGC(Ala)_ and four tRNA^Ala^ variants containing synonymous anticodons decoding leucine codons. Proportion of mistranslated peptides at each codon was calculated from mass spectrometry data as the number of unique mistranslated peptides divided by the number of wild-type peptides where the leucine residue is coded for by the codon indicated. Only peptides containing one leucine residue were considered. Each point represents one biological replicate (n ≥ 3). **(B)** Venn diagrams showing the overlap between the unique peptides (left) and proteins (right) where mistranslation was detected by mass spectrometry in strains containing tRNA^Ala^ variants that have one of four synonymous anticodons decoding leucine codons. **(C)** Codon specific proportion of unique mistranslated peptides for the control strain containing tRNA^Ala^ _GGC(Ala)_ and four tRNA^Ala^ variants containing synonymous anticodons decoding valine codons as in (A). **(D)** Venn diagrams showing the overlap between the unique peptides (left) and proteins (right) where mistranslation was detected by mass spectrometry in strains containing tRNA^Ala^ variants that have one of four synonymous anticodons decoding valine codons.

The same analysis was done for the set of tRNA^Ala^ variants substituting alanine for valine. The estimated total mistranslation by variants with UAC(Val) and CAC(Val) anticodons was similar but these variants decreased growth to 87% and 54% of the control strain, respectively. The decoding specificity of the variants with UAC(Val) and CAC(Val) anticodons was distinct from each other and from the variants with AAC(Val) and GAC(Val) anticodons, which were similar (Figure 6C). Again, in line with the differences in decoding specificity, there was minimal overlap in the mistranslated peptides and proteins identified in strains containing the variants with UAC(Val) and CAC(Val) anticodons (Figure 6D), while variants with AAC(Val) and GAC(Val) anticodons mistranslated relatively similar sets of peptides and proteins. These results suggest that differences in decoding specificity could contribute to the impact on growth of synonymous anticodon variants.

### Decoding specificity of tRNA^Ala^ variants

The mass spectrometry data identifying sites of mistranslation allowed us to determine the codons mistranslated by each variant. Figure 7 shows the decoding specificity for tRNA^Ala^ variants with statistically elevated mistranslation frequencies (raw data in Table S9, Figure S5). In 10 of 13 cases, G at position 34 resulted in mistranslation of codons with C and U at the third codon position. Interestingly, in these cases, with the exception of GCC(Gly), a greater proportion of mistranslated peptides were identified with codons ending in a 3’ C rather than a 3’ U. Variants with A at position 34 have the potential to be modified to I34 (inosine) and decode codons ending with U, C and A [58–60]. Of the nine A34-containing variants, seven mistranslated codons ending in U and C, with the exceptions being ACC(Gly) and AUG(His). This suggests these variants have expanded decoding consistent with inosine modification; however, we only detected mistranslation at codons ending in A for the AGG(Pro) variant. Variants with U at position 34 predominantly mistranslated codons with 3’ A, with U:G decoding only detected for the UCG(Arg) variant. Interestingly, U:U decoding, indicative of superwobble, was detected for UGG(Pro) and UAC(Val) variants. Mistranslation by tRNA^Ala^ variants with C at position 34 was limited to codons ending in G.

**Figure 7.**
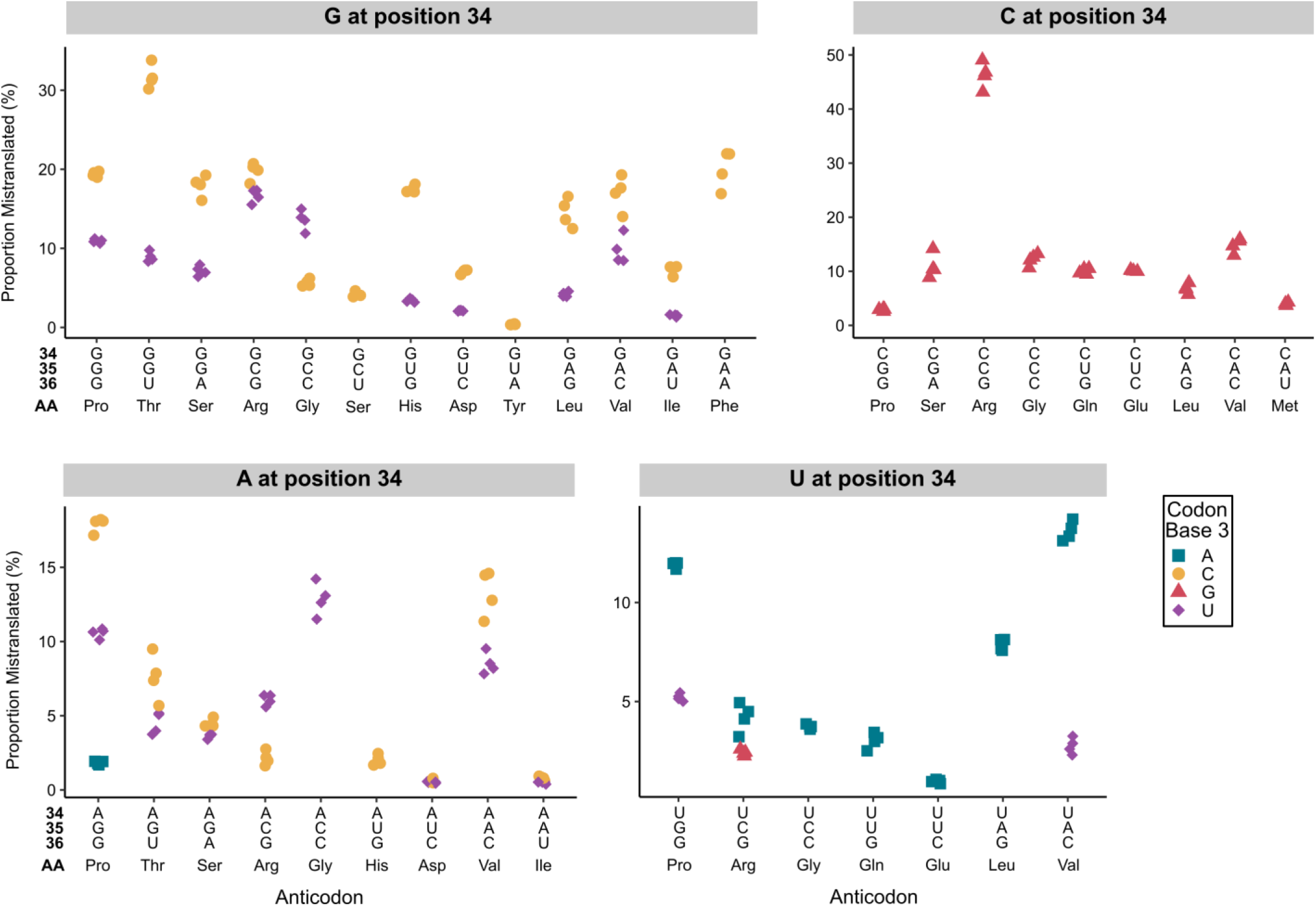
Profile of mistranslation events detected at synonymous codons for tRNA^Ala^ variants with G, A, C or U at base 34. For each tRNA^Ala^ anticodon variant, the proportion of mistranslated peptides identified at each synonymous codon was calculated from the number of unique peptides with a mistranslation event at that codon relative to the number of unique peptides containing a wild-type residue at that specific codon. In all cases, positions 35 and 36 formed Watson-Crick pairs with positions 2 and 1 of the codon, respectively. To confidently localize the mistranslation event and identify the mistranslated codon, only peptides containing a single target amino acid were used in this analysis. Only mistranslation that is statistically above the frequency measured for the same codon in the wild-type strain is shown (Welch’s *t*-test; Benjamini-Hochberg corrected *p* < 0.01). Each point represents one biological replicate (n ≥ 3).

## Discussion

Our study has characterized the 60 non-alanine tRNA^Ala^ anticodon variants for their mistranslation potential and impact on the growth of *Saccharomyces cerevisiae*. Twenty-five of the 60 variants reduced growth when present in single copy. Slow growth due to some but not all remaining variants was observed when their copy number was increased. These results suggest that any potential impact of tRNA^Ala^ anticodon variants on human health or disease will depend on the specific sequence of the anticodon. Mistranslation was observed for 52 of 57 non-alanine variants, indicating that mistranslation does not always lead to reduced growth. The 52 variants that do mistranslate will have applications in synthetic biology and could potentially be used as therapeutics. Furthermore, the fact that these tRNA^Ala^ variants can be introduced into cells without causing cell death has implications for the adaptability and evolution of the genetic code (see for example [61,62]).

### Factors influencing the impact of tRNA^Ala^ variants on cell growth

There are multiple factors that interface to influence the impact of mistranslating tRNAs on cell growth. The first to consider is the overall level of mistranslation. The amount of mistranslation experienced by a cell is dependent on the frequency of mistranslation, the number of codons that are mistranslated and the expression level of proteins containing the effected codons. Kinetic aspects of translation are relatively uniform for native aminoacyl-tRNAs despite the diversity in amino acid structure and tRNA sequence [63–66]. At first glance, this would suggest that all tRNA^Ala^ anticodon variants should mistranslate at similar levels and therefore mistranslation frequency would only depend on the number of competing native tRNAs. However, as noted by the Uhlenbeck lab [63,67], to obtain translational uniformity each tRNA has evolved sequence or modification differences to tune their activity to a similar level. Our results support this argument since when taken out of their native context, the tRNA^Ala^ anticodon variants do not elicit an equivalent level of mistranslation or impact on growth. This is most easily seen for 2-box anticodons where there is only one native isodecoder in *S. cerevisiae* (Phe, Asn, Asp, His, Tyr and Cys), and thus the competition with native tRNAs is the same for both synonymous anticodons. For all six of these amino acid families, the two tRNA^Ala^ variants have different estimated mistranslation frequencies and different impacts on growth (Figure S6). In all of these cases, the variant containing a G at position 34 had a more severe impact than the A-containing counterpart. In fact, throughout the entire set of variants G/C-rich anticodons had a greater impact on growth compared to the A/U-rich variants. It is possible that tRNA modifications play a role in this effect by tuning translation efficiency. For example, native tRNAs with A/U rich anticodons are often modified at base 37 to N^6^ -isopentenyladenosine or N^6^ -threonylcarbamoyladenosine to stabilize base pairing interactions [68–71]. These modifications likely do not occur for tRNA^Ala^, which has a G/C rich anticodon, thus potentially reducing the activity of A/U rich anticodon variants. Consistent with this, Pernod *et al*. found limited codon:anticodon interactions between A/U rich codons in an *in vitro* system [72].

A second principal factor determining the impact of each mistranslating tRNA^Ala^ variant is the specific set of proteins and peptides that are mistranslated. Even small changes in expression or function of certain proteins can impact growth [73]. Since amino acid composition varies from protein to protein, each tRNA^Ala^ anticodon variant has the potential to contribute to slow growth by decreasing the functional amount of one or more distinct proteins. The ability of variants to impact different cellular pathways is supported by our previous finding where two tRNAs mistranslating at similar frequencies, one mis-incorporating alanine at proline codons and the other serine at arginine codons, had unique sets of genetic interactions [74]. Furthermore, because codon usage is not random, synonymous anticodons target different proteins and thus impact growth to different extents as demonstrated by tRNA^Ala^ variants decoding the variants with UAG(Leu) and CAG(Leu) anticodons. Also related to mistranslation contributing to changes in protein expression is the ability of specific mistranslation events to influence translational elongation and regulate protein folding. Reduced translation rate at the 5’ end of genes, which minimizes abortive synthesis, and ribosome stalling, which can promote protein folding, are achieved by the selection of specific codons (reviewed in [75–77]). Proline codons specifically have a significant role since peptide bond formation involving proline as a donor or acceptor is slow and stretches of two or more prolines lead to ribosome stalling. Therefore, mistranslation events replacing proline have the potential to alter protein expression. In this regard, it is interesting that tRNA^Ala^ variants with proline anticodons are, as a group, the most impactful on growth and the only set where a positive correlation between the amount of mistranslation and growth was observed.

Related to the importance of codon usage regulating protein expression, Gamble *et al*. identified 17 codon pairs in yeast that negatively impact translation [78]. These codon pairs tend to be conserved throughout yeasts [79], suggesting they have a functional role. When non-native tRNAs with anticodons cognate to one of the codons in the inhibitory pair are introduced into cells, the inhibitory effect of the pair on translation is alleviated [78]. Likewise, introducing the non-native mistranslating tRNA^Ala^ variants might affect the regulation of gene expression achieved through inhibitory codon pairs. Interestingly, three of the 17 codon pairs contain the proline CCG codon, providing another possible explanation for why the CGG(Pro) anticodon variant had the largest impact on growth while having relatively low levels of mistranslation.

We also considered the nature of the amino acid substitution as a determinant of the impact of each tRNA^Ala^ variant on growth. The methyl side chain of alanine is small and hydrophobic. Mis-incorporating alanine for amino acids with distinct chemical or functional properties might be expected to have a greater impact. Supporting this argument, the correlation between the impact on growth and abundance of mistranslated peptides improved when the anticodons were separated by amino acid substitution. When we considered five properties that differentiate the 20 amino acids [57], differences in volume, electron-ion interaction potential and π-helix propensity of the replaced amino acid relative to alanine contributed to the impact each substitution had on growth.

In this discussion we have restricted our consideration to the roles of tRNAs in translation. However, many reports are continuing to expand the known roles for tRNAs in other processes. For example, tRNA^Gln^ _CUG_ is important for signaling in response to poor nitrogen sources [80] and the anticodon is required to impart this function [81]. We can not exclude the possibility that these non-translational functions contribute to the growth impairment of some of the tRNA^Ala^ variants.

### Similarities and differences with mistranslating tRNA^Ser^ variants

Previously, Zimmerman *et al*. investigated the impact of tRNA^Ser^ variants with all 58 non-serine anticodons on the growth of yeast [13]. While their data was acquired in pooled format, making direct comparison with our tRNA^Ala^ results difficult, their findings of anticodon specific differences including those between synonymous anticodons are consistent with our tRNA^Ala^ results.

We previously analyzed some individual tRNA^Ser^ variants [14] and find that in general, tRNA^Ser^ anticodon variants are more toxic than their tRNA^Ala^ counterparts. Mistranslation by a tRNA^Ala^ or tRNA^Ser^ variant would be reduced if its aminoacylation by AlaRS or SerRS, respectively, was in competition with the anticodon-cognate aaRS. This neutralizing mechanism would be most likely for the 11 aaRSs that use the full anticodon as a primary identity element for aminoacylation (Asn, Asp, Cys, Gln, His, Ile, Lys, Met, Phe, Thr and Trp) [17] and for native anticodon sequences. Anticodon cognate aminoacylation may account for differences between tRNA^Ala^ and tRNA^Ser^ variants since the unique variable arm of tRNA^Ser^ is likely to negate interaction with the anticodon cognate aaRSs.

### Decoding specificity of anticodon variants

In addition to decoding their exact Watson-Crick base pairing codons, expanded decoding by tRNAs occurs through G:U wobble pairing and modification of base 34 [82]. Our dataset allowed us to investigate the *in vivo* decoding specificities for mistranslating tRNA^Ala^ anticodon variants. For most variants with G, C or U at position 34, the Watson-Crick cognate codon was mistranslated the most frequently. Variants with A34 mistranslated codons ending in C and U and to a lesser extent A, consistent with A34 being modified to inosine [58,83]. The exception was the ACC(Gly) anticodon variant, which is not native in yeast. Interestingly, we only detected mistranslation at codons ending in A for the AGG(Pro) variant. The need for mechanisms to regulate decoding by variants modified to have I34 may reflect that for two-box tRNA families, I:A decoding would result in translation across the box. It is possible that I:A decoding is minimized in non-cognate circumstances to prevent across the box mistranslation should the A34 arise through mutation of native two-box tRNAs. Consistent with this, we did not observe across the box alanine mis-incorporation for two-box anticodon variants with A34.

Anticodon variants with G or U at position 34 also exhibited expanded decoding. For 10 of 13 variants with G at position 34, codons ending in U were mistranslated. Conversely, only the UCG(Arg) variant decoded codons ending in G. This agrees with previous reports that G:U (anticodon:codon) pairs are more favorable than U:G pairs, unless U is modified [56,72,84,85]. For two variants with U at position 34 (UGG(Pro) and UAC(Val) anticodons), we observed decoding of codons ending in U. This suggests ‘superwobble’ where anticodons with U at position 34 decode codons ending in all four nucleotides [86] and is consistent with the ability of *S. cerevisiae* tRNA^Pro^ _UGG_ to decode all four proline codons [87].

## Data availability

Raw images and data are available in the supplemental information. The raw mass spectrometry data have been deposited to the ProteomeXchange Consortium via the PRIDE[51] partner repository with the dataset identifier PXD038242 and an annotated list of the file names can be found in Supplemental Table S4. The mass spectrometry data can be reviewed using username: and password: YIi3WK2a.

## Author contributions

E.C, C.J.B. and M.D.B conceived and developed the project. E.C. and J.G. cloned the tRNA variants. E.C. conducted the growth experiments, analyzed the data, identified factors correlating with mistranslation abundance and growth impact and created figures 1, 2, 3 and 4 and associated supplemental figures. M.D.B conducted the mass spectrometry experiments with assistance from R.A.R.-M., analyzed the data and created figures 5, 6 and 7 and associated supplemental figures. M.R. constructed multicopy variants. M.R. and M.D. participated in data analysis. C.J.B supervised the work and provided funding. J.V. supervised the mass spectrometry work and provided funding. M.D.B. supervised the project. E.C., C.J.B and M.D.B wrote the manuscript and all authors edited the manuscript.

## Acknowledgments

We would like to thank Josh Isaacson and Josephine Davey-Young for critically reading the manuscript and Tallulah Andrews and members of the Villén lab for insightful discussions.

