## Supplemental Figures for "Anticodon sequence determines the impact of mistranslating tRNA^Ala^ variants"

AAGCTTC GGACGATTGC CAACCGCCGA AAAGGTTCAA GCCAAGAACA AAAAGTAGAG<br>
AGAATACCCC AACAATAGCA TTTGATGGCT AAACGGCAGT GCCCGTTAAT ACTATTCAAA<br>
TTATACCCCG CATTAATAAT TCCTGTCCCT TTAGTCTCGT CCTTCAGCCG TATAGCCGCA<br>
AAACTTCGTT CAATGATTTC ATGCCATCAT TACACCGCGA GTCGTGGGTG CATTGTAGTT<br>
TATTACGAAA AGTATCTACA ATACTTGCTT AAATAACCTA CATTGTTTTA **GGGCGTGTGG**<br>
**CGTAGTCGGT AGCGCGCTCC CTTAGCATGG GAGAGGTCTC CGGTTCGATT CCGGACTCGT**<br>
**CCA**TTATTTT TTTATTTTTA TTTTTTTTTT ATCGCTTACT GATTATCAGA TATCTTCGAC<br>
AGCCTCACAC AGATTGGTGA TCACGCACCC ATAATCATTT CTTCGGGCAT GCTCCTATTA<br>
GCCACGGTTC GCAGAATAAT CTGCGCGATG TTATTCACCA AGACCGTGGA GTCTCCTTTT<br>
CGGTGGGATC CGCTTATATC CGTATGCGTC GGTTGCTTTC TCGGGGAAAG GAAAAGGAGA<br>
AAGCCCGGAT AACACGACAC AAGAGTCATT CGTTATGCAC GGACACAGGG AATCGTGCGG<br>
AGCCGGGGAT AAGAGGCCGT GCTAATTTCC GCGGAAGCGG CCGC

**Figure S1. Inducible alanine mistranslating tRNA with tRNA$^{Ser}$ *SUP17* flanking sequence.** Red bolded sequence represents the alanine tRNA. Underlined sequence is the restriction sites used to clone the construct into the yeast plasmid.

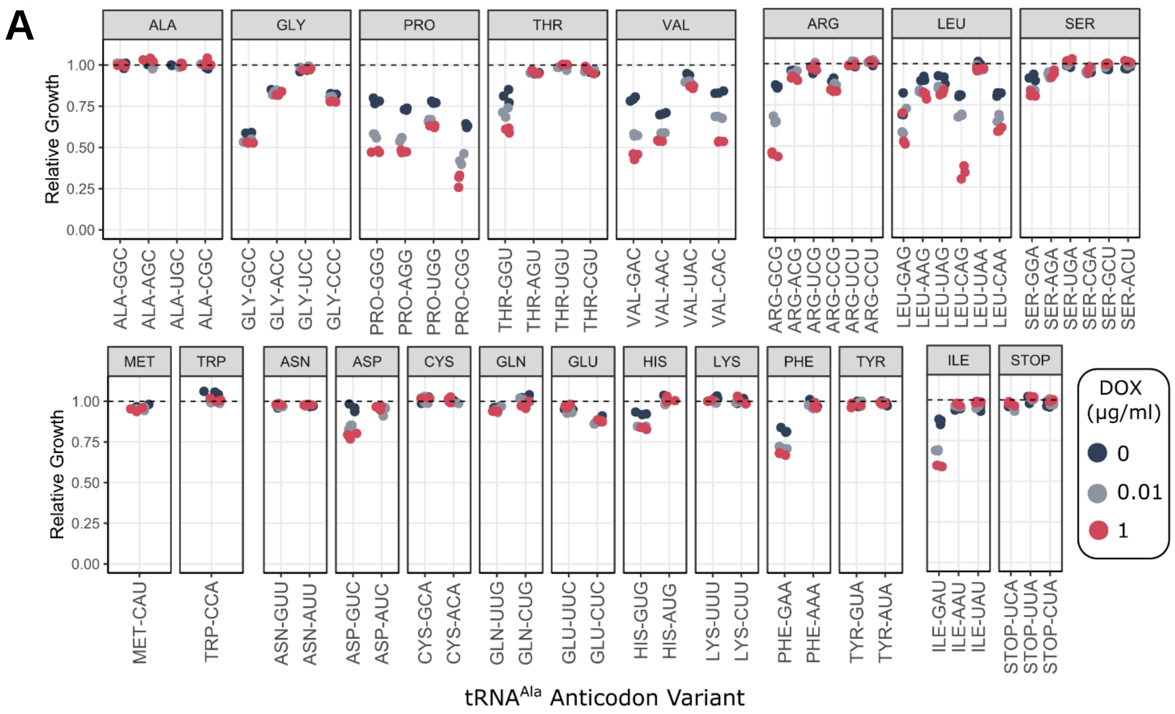


**Figure S2. Growth impact of tRNA$^{Ala}$ anticodon variants at different doxycycline concentrations.** Relative growth of all 64 tRNA$^{Ala}$ anticodon variants compared to control tRNA$^{Ala}_{GGC(Ala)}$ at 0, 0.01 and 1.0 µg/mL doxycycline represented by the blue, grey, and red points, respectively. Each point is a biological replicate (n = 3). Plasmids containing each of the 64 variants were transformed into strain CY8652. Cultures were grown in triplicate for 48 hours at 30°C in 2% glucose medium lacking uracil and leucine, diluted to an $OD_{600}$ of 0.1 in the same medium with 0, 0.01 or 1.0 µg/mL doxycycline and grown for 24 hours at 30°C with agitation. $OD_{600}$ was measured every 15 minutes and doubling time was quantified with R package 'growthcurver' and used to calculate relative growth.

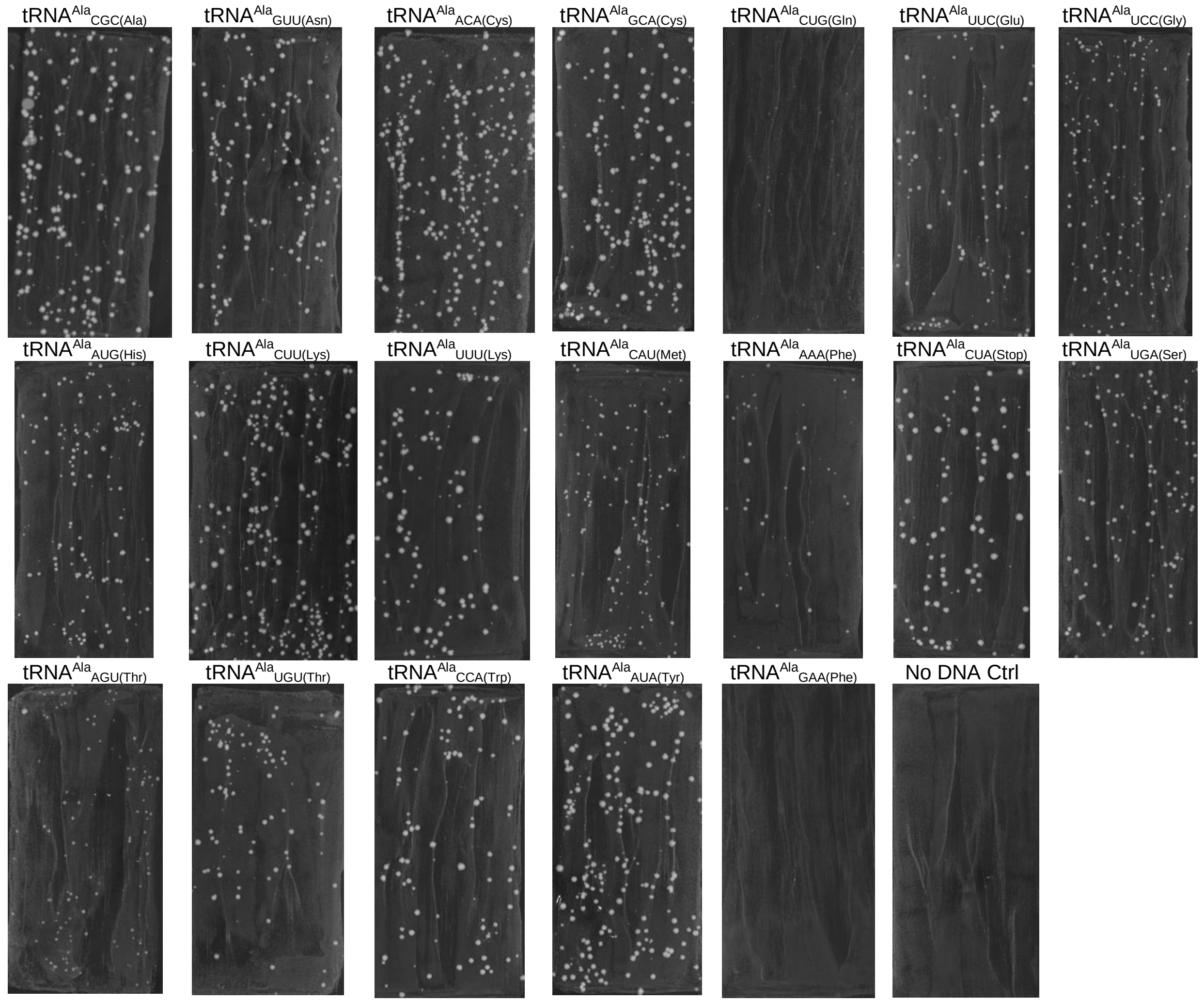


**Figure S3. Multicopy $tRNA^{Ala}$ anticodon variant transformants.** Strain BY4742 was transformed with 1.0 µg of each plasmid containing the indicated variant, plated on medium lacking uracil and grown at 30°C. Transformants were imaged after 48 hours. $tRNA^{Ala}_{GAA(Phe)}$ is a control as it mistranslates when expressed from a centromeric plasmid.

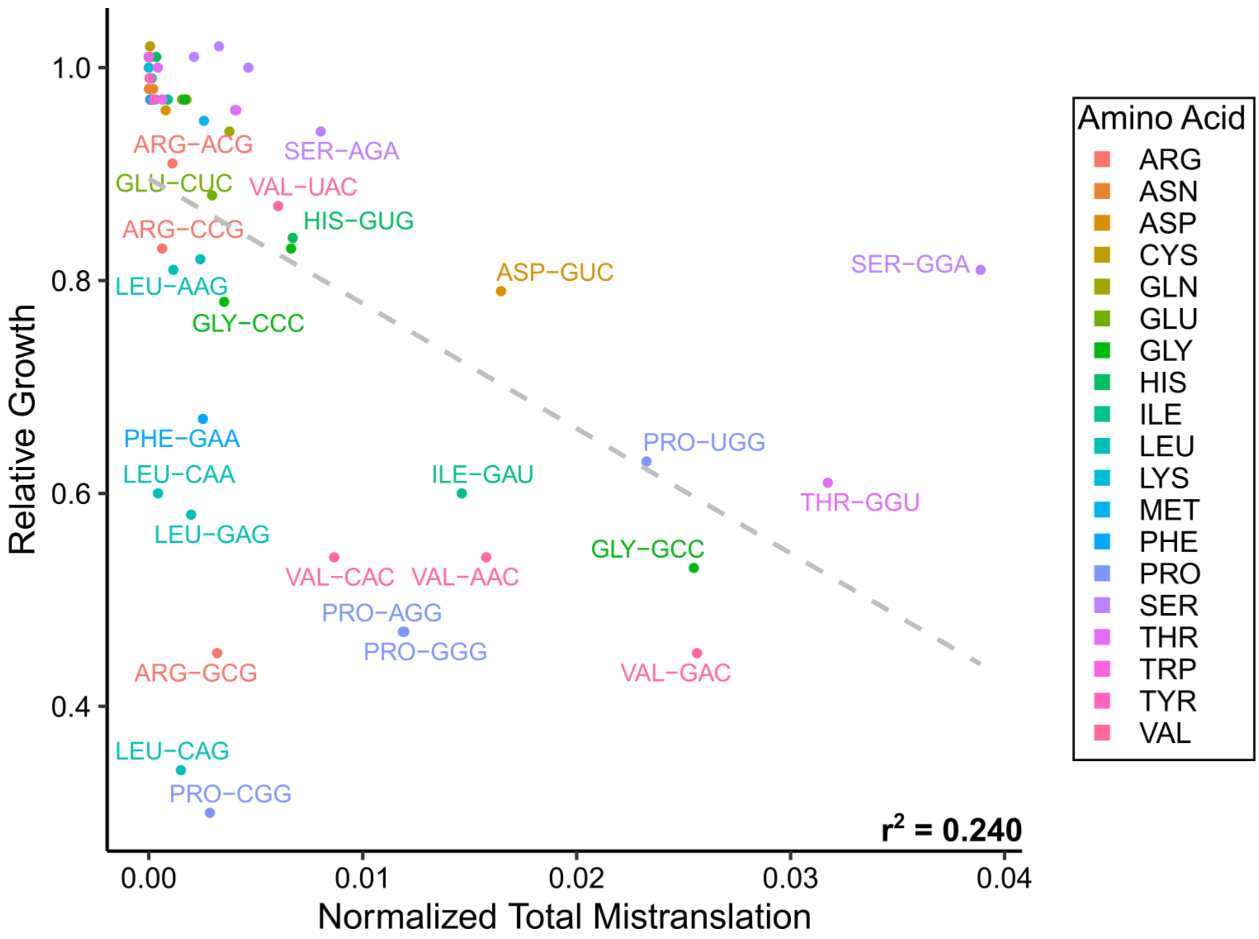


**Figure S4.** Summed MS1 intensity of mistranslated peptides normalized to the summed MS1 intensities of all detected peptides (normalized mistranslation abundance) plotted against the relative growth for each variant. The grey dashed line represents the correlation between the two variables. The linear $r^2$ value is shown.

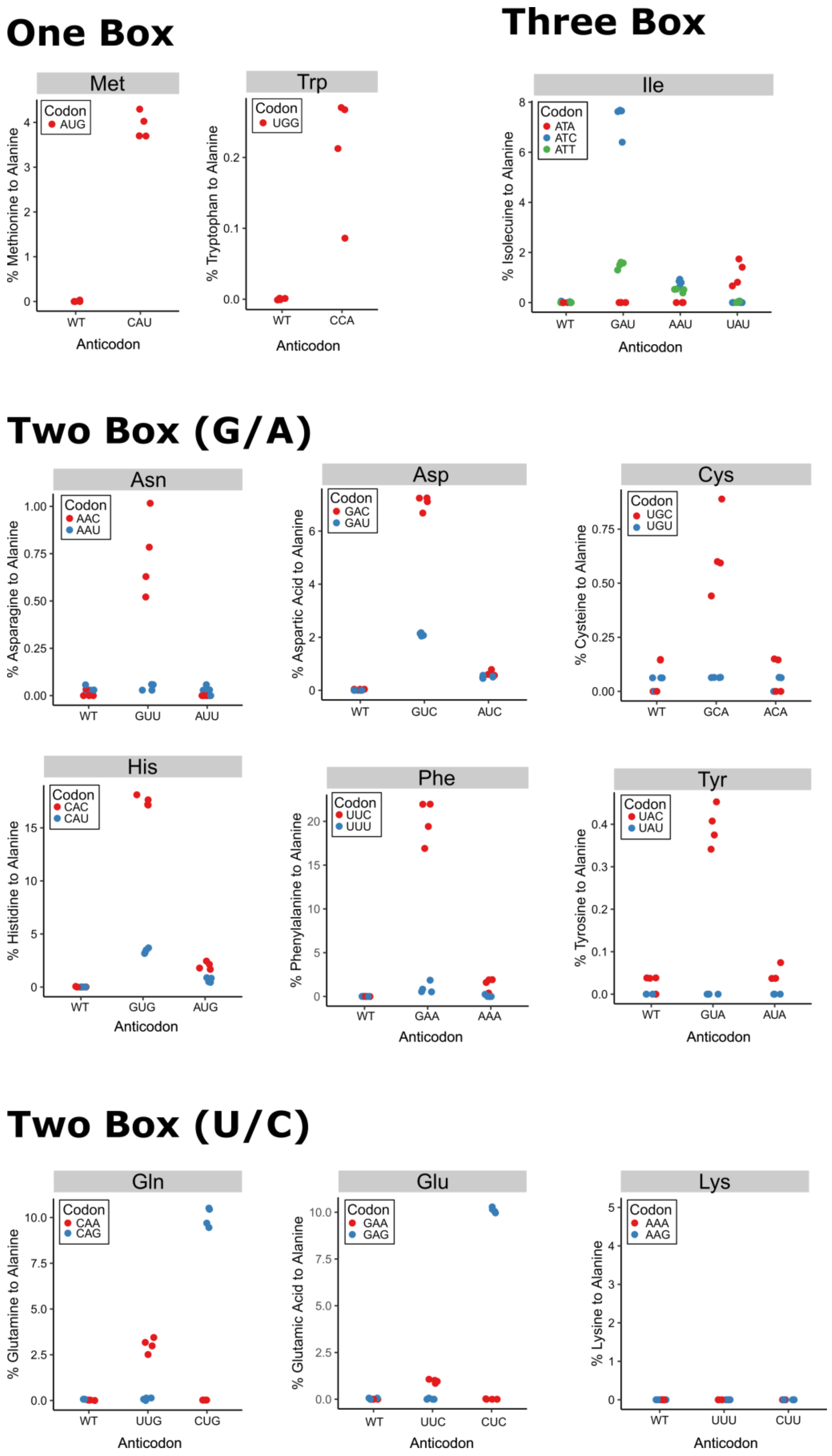
One Box
Three Box
Met
Codon
AUG
% Methionine to Alanine
0
1
2
3
4
WT
CAU
Anticodon
Trp
Codon
UGG
% Tryptophan to Alanine
0.0
0.1
0.2
WT
CCA
Anticodon
Ile
Codon
ATA
ATC
ATT
% Isolecuine to Alanine
0
2
4
6
8
WT
GAU
AAU
UAU
Anticodon
Two Box (G/A)
Asn
Codon
AAC
AAU
% Asparagine to Alanine
0.00
0.25
0.50
0.75
1.00
WT
GUU
AUU
Asp
Codon
GAC
GAU
% Aspartic Acid to Alanine
0
2
4
6
WT
GUC
AUC
Cys
Codon
UGC
UGU
% Cysteine to Alanine
0.00
0.25
0.50
0.75
WT
GCA
ACA
His
Codon
CAC
CAU
% Histidine to Alanine
0
5
10
15
WT
GUG
AUG
Anticodon
Phe
Codon
UUC
UUU
% Phenylalanine to Alanine
0
5
10
15
20
WT
GAA
AAA
Anticodon
Tyr
Codon
UAC
UAU
% Tyrosine to Alanine
0.0
0.1
0.2
0.3
0.4
WT
GUA
AUA
Anticodon
Two Box (U/C)
Gln
Codon
CAA
CAG
% Glutamine to Alanine
0.0
2.5
5.0
7.5
10.0
WT
UUG
CUG
Anticodon
Glu
Codon
GAA
GAG
% Glutamic Acid to Alanine
0.0
2.5
5.0
7.5
10.0
WT
UUC
CUC
Anticodon
Lys
Codon
AAA
AAG
% Lysine to Alanine
0
1
2
3
4
5
WT
UUU
CUU
Anticodon

### Four Box

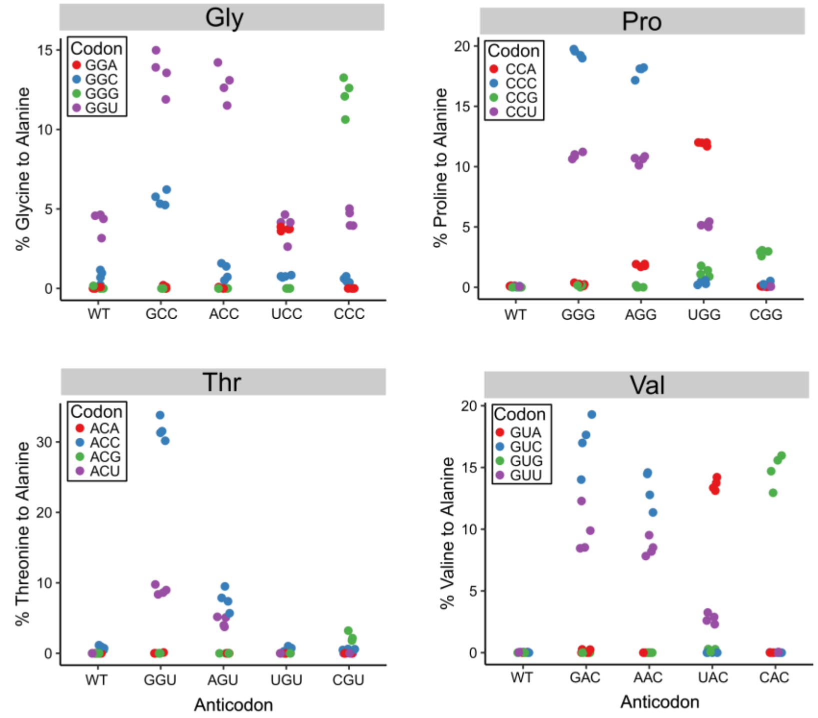


### Six Box

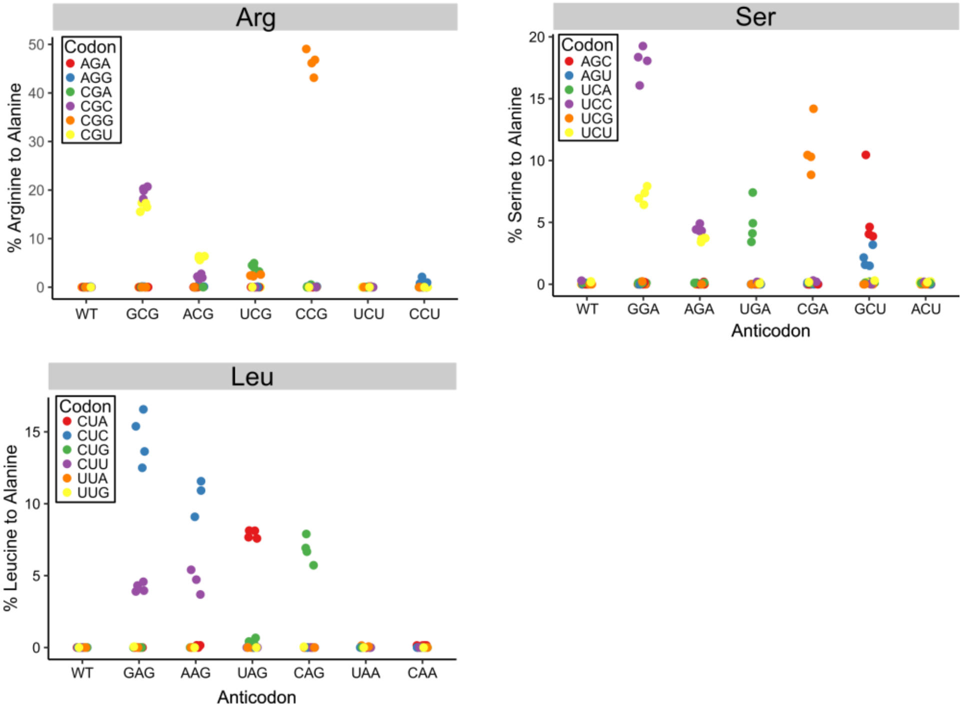

**Figure S5. Proportion of mistranslated peptides identified per codon for the tRNA$^{Ala}$ variants.** Yeast strain CY8652 containing either a plasmid with a control tRNA$^{Ala}_{GGC(Ala)}$ or a plasmid containing an tRNA$^{Ala}$ anticodon variant were grown in medium containing 1.0 µg/ml doxycycline to induce tRNA expression and harvested at an $OD_{600}$ of ~ 1.0. Mass spectrometry analysis of the cellular proteome was performed after proteins were extracted in a denaturing buffer. Mistranslation frequency at all synonymous codons for each variant was calculated from the number of unique peptides where alanine mis-incorporation was observed at a specific codon relative to the number of unique peptides observed where the wild-type amino acid was present at the same codon. Only peptides containing a single target residue were used. Each point represents one biological replicate ($n \geq 3$).

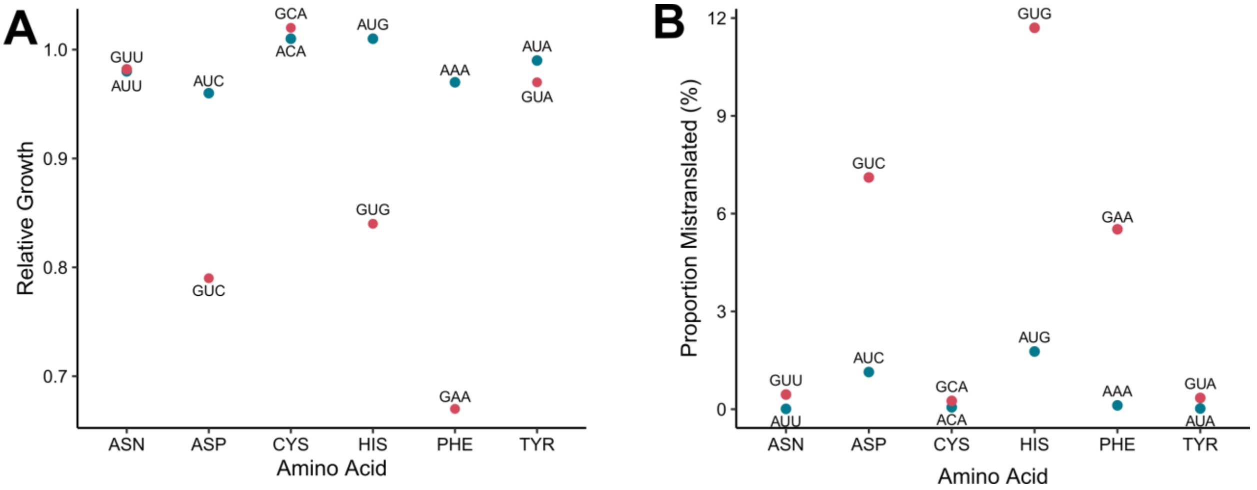


**Figure S6.** Average relative growth (A) or proportion of peptides identified as mistranslated (B) is shown for six different amino acids where only one of the two anticodons naturally exists in yeast (i.e. the tRNA$^{Ala}$ anticodon variants are both competing to mistranslate against the same set of native tRNAs).
